# Odor mixture perception: can molecular complexity be a factor determining elemental or configural perception?

**DOI:** 10.1101/2020.05.27.120154

**Authors:** Masayuki Hamakawa, Hiroya Ishikawa, Yumika Kikuchi, Kaori Tamura, Tsuyoshi Okamoto

## Abstract

Odor mixtures can evoke smells that differ from those of their individual odor components. Research has revealed the existence of two perceptual modes, in which a mixture can be perceived as either the original smells of its individual components (elemental) or as a novel smell (configural). However, the factors underlying the perceptual transformation that occurs when smelling a mixture versus its original components remain unclear. Therefore, the present study aimed to identify the properties of odorants that affect olfactory perception of odor mixtures, focusing on the structural complexity of an odorant. We conducted psychophysical experiments in which different groups of participants were instructed to provide olfactory perceptual descriptions of low-, medium-, and high-complexity odor mixtures or components, respectively. To investigate the perceptual modes induced by the mixtures, we compared the participants’ evaluations between mixtures and components via two types of analyses. First, we compared each olfactory description following quantification via principal component analysis. We then compared data based on seven major olfactory perceptual groups. We observed that odor mixtures composed of low-complexity odorants were perceived as relatively novel smells with regard to both minor (olfactory descriptions) and major (perceptual community) odor qualities than medium- and high-complexity mixtures. Such information may further our understanding of the olfactory perceptual modes of odor mixtures.

## Introduction

Odor mixtures can evoke smells that differ from those of their individual odor components. Various research groups have proposed the existence of two different olfactory perceptual modes: elemental and configural^1–6^. In the elemental mode, a mixture is perceived as the original smells of its components, whereas in the configural mode, a mixture is perceived as a new smell. Previous studies have demonstrated that these perceptual modes occur not only alternatively, but also simultaneously^1–3,5,7^. As for the factors determining the perceptual modes (elemental or configural), several challenges have been conducted to elucidate the factors. Some studies have suggested that the capacity to identify the odor components of a mixture depends on the number of components^3,5,7,8^, and others have suggested that the perceptual mode is determined based on an individual’s olfactory background (*e.g.*, olfactory learning and pre-exposure)^1,5,9^. A recent research indicated that specific odorants, as terms “key odorants”, can induce the elemental perception, and specific associations of odorants, as terms “key association”, can do the configural perception^10^.

An important first step towards understanding how the perceptual mode of an odor mixture is determined would be to focus on molecular properties of a single odorant in the mixture. Research has indicated that process of olfactory perception is initiated by neural coding in the peripheral olfactory system via olfactory receptor neurons^11,12^. Information from the receptors is then sent to the olfactory bulb and eventually terminates in olfactory-related areas in the cerebral cortex^13–15^. Previous studies have suggested that the pattern of neural activity in each stage of olfactory processing depends on the odorant molecular feature (*i.e.*, an odorant’s carbon structure and functional group)^13,16^. Such odorant molecular feature can affect also olfactory perception^17–19^. Such findings imply that certain aspects of an odorant molecule determine how it is perceived. However, it remains unclear how odorant characteristics reflecting multiple molecular features play a role in determining the perceptual mode.

Molecular complexity is characterized by several structural features such as bond connectivity, symmetry, and atomic components^20^. Although some recent studies have reported that the complexity of monomolecular odorants indeed affects olfactory perception^21,22^, to our knowledge, no studies have examined the relationship between the perceptual mode of odor mixtures and molecular complexity.

In the present study, we aimed to clarify whether the complexity of odorant molecules influences the perceptual mode of odor mixtures. A previous study suggested that according to the odorant molecular structural complexity, the number of activated olfactory receptors was varied which leads to the change of the variety in evoked olfactory perceptual descriptors. We therefore hypothesized that according to the molecular complexity, the degree of the perceptual modes (elemental or configural) is determined (*e.g.*, mixtures composed of low-complexity odorants is relatively perceived as configural than high-complexity mixtures). To test it, we prepared 12 odor components and 18 binary odor mixtures, which were divided into three groups according to their molecular complexity scores based on a previous study^21^ (low, medium, and high). We then conducted a psychophysical experiment wherein naïve participants were instructed to describe the smells of the components or mixtures by selecting from among several linguistic expressions (referred to as olfactory notes: *e.g.*, “woody” and “citrusy”). Each participant was also asked to rank the selected olfactory notes based on the relative intensity of each perceived smell. We then examined differences in the relative intensity of olfactory notes between odor mixture and its component odors using two types of analyses: comparison of individual olfactory notes quantified by principal component analysis; comparison based on major olfactory perceptual groups established in a previous study^23^. Our analysis suggested that molecular complexity plays a role in determining the perceptual mode of odor mixtures.

## Experimental

### Participants

Fourteen healthy Kyushu university students (eight women and six men, mean age ± standard error of the mean = 22.1 ± 0.57 years) participated in the odor mixture experiment, while an additional ten healthy students (two women and eight men, mean age ± standard error of the mean = 21.9 ± 0.877 years) participated in the odor component experiment. Participants of the two experiments did not overlap. All participants reported having normal olfaction, none reported a history of psychiatric disorders, and no history of special olfactory training. No participants were excluded from the data analysis. The ethics committee of the Faculty of Arts and Science at Kyushu University approved all experimental stimuli, protocols, and procedures (201510R2). Written informed consent was obtained from each participant. All methods of this research were performed in accordance with the approved guidelines.

### Olfactory stimuli

The molecular complexity values of the odorants were obtained from the PubChem database of chemical molecules (https://pubchem.ncbi.nlm.nih.gov). The molecular complexity of an odorant in the database was calculated based on its structure, including its bond connectivity, diversity of non-hydrogen atoms, and symmetry^20^.

A total of 12 odor components were used in the present study (see Table 1 for information regarding concentration, solvent, and associated olfactory notes). The 12 odor components were selected as follows: (1) Odorants with only two olfactory notes listed in a commercial-release database, *Sigma Aldrich Ingredients Catalog: Flavors & Fragrances* (http://www.sigmaaldrich.com), were extracted from the 128-odorant collection published in a previous report^24^; and (2) the extracted odorants were sorted according to their molecular complexity. Note that methylsulfanylmethane (Chemical Abstracts Service Number = 75-23-8, complexity = 2.8) and decanoic acid (Chemical Abstracts Service Number = 334-48-5, complexity = 110) were removed because the former was deemed hazardous to the participants, and the latter could not be detected via gas chromatography/mass spectroscopy (GC/MS) analysis. The concentration of all odor components for preparation of the 18 mixtures, as shown in Table 1, was determined based on the 128-odorant collection, in which the odor intensity of individual component was almost equivalent^24^. Olfactory notes used in the present study were obtained from the Sigma Aldrich database, although participants were also permitted to provide their own terms.

**Table 1.**
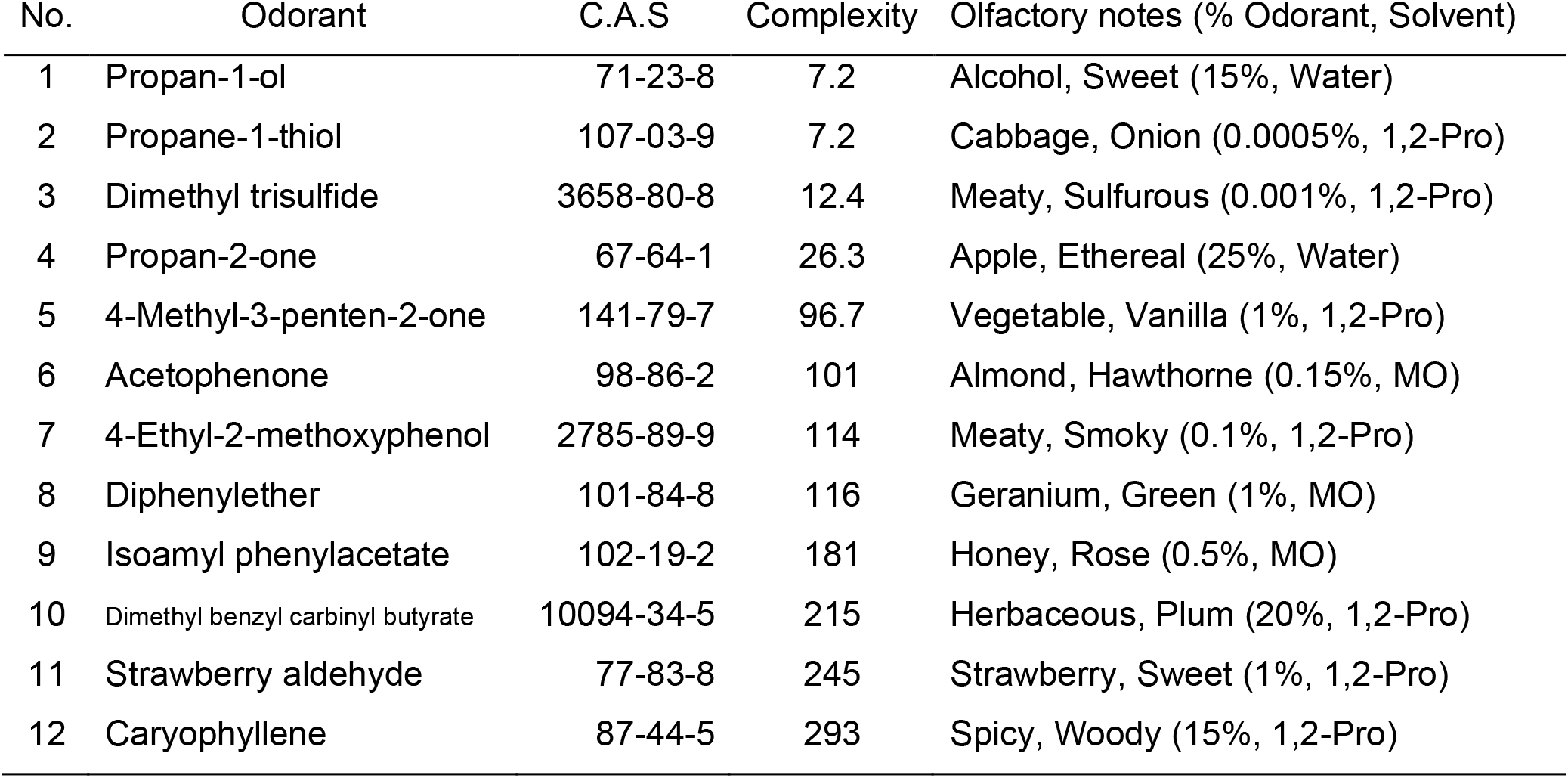
Odor components and their associated properties and solvent conditions. C.A.S. is Chemical Abstracts Service Number. 1,2-Pro is 1,2-Propanediol. MO is mineral oil.

In order to simply test our hypothesis: the degree of the perceptual mode is determined according to the molecular complexity, we categorized the selected odorants (ranges given in parentheses) into the following three groups based on molecular complexity values: The four lowest complexity odorants were classified into the low group (7.2-26.3), the four highest complexity odorants were classified into the high group (181-293), and the four odorants with complexities of approximately 100 were categorized into the medium group (96.7-116). This manner of classification is consistent with methods reported in a previous study^21^. Although there were the 66 possible binary combinations from the 12 components, the four odorants in each complexity category were combined to create six binary odorant mixtures (18 mixtures in total) in this study. All odor mixtures used in the present study are listed in Table 2. The complexity score of an odor mixture was defined as the sum of its binary components.

**Table 2.** Odor mixtures used in the present study.

| No. | Component 1 | Component 2 | Sum of complexity |
| --- | --- | --- | --- |
| 1 | Propan-1-ol | Propane-1-thiol | 14.4 |
| 2 | Propan-1-ol | Dimethyl trisulfide | 19.6 |
| 3 | Propane-1-thiol | Dimethyl trisulfide | 19.6 |
| 4 | Propan-1-ol | Propan-2-one | 33.5 |
| 5 | Propane-1-thiol | Propan-2-one | 33.5 |
| 6 | Dimethyl trisulfide | Propan-2-one | 38.7 |
| 7 | 4-Methyl-3-penten-2-one | Acetophenone | 197.7 |
| 8 | 4-Methyl-3-penten-2-one | 4-Ethyl-2-methoxyphenol | 210.7 |
| 9 | 4-Methyl-3-penten-2-one | Diphenylether | 212.7 |
| 10 | Acetophenone | 4-Ethyl-2-methoxyphenol | 215 |
| 11 | Acetophenone | Diphenylether | 217 |
| 12 | 4-Ethyl-2-methoxyphenol | Diphenylether | 230 |
| 13 | Isoamyl phenylacetate | Dimethyl benzyl carbiny l butyrate | 396 |
| 14 | Isoamyl phenylacetate | Strawberry aldehyde | 426 |
| 15 | Dimethyl benzyl carbiny l butyrate | Strawberry aldehyde | 460 |
| 16 | Isoamyl phenylacetate | Caryophyllene | 474 |
| 17 | Dimethyl benzyl carbiny l butyrate | Caryophyllene | 508 |
| 18 | Strawberry aldehyde | Caryophyllene | 538 |

We prepared the mixtures as follows: A total of 5.0 μL of each of the two component solutions was pipetted onto a cotton cloth (1 cm x 1 cm), following which the two cloths were placed together in a sealed 20-mL vial at room temperature (set to 25°C using an air-conditioner) for 5 min. After 5 min, the cotton cloths were removed, and the vials containing the gaseous odor mixtures were presented to each participant. The preparation of individual odor components was same to the mixtures.

### Experimental procedures

Two psychophysical experiments were performed: an odor mixture experiment and an odor component experiment.

The odor mixture experiment was conducted in a well-ventilated room over the course of 2 days, with sessions separated by 24 h. The order in which the 18 odor mixtures were presented was randomized (computer-generated) from day to day for each participant. During the evaluation of each mixture, participants were instructed to open the vial containing the mixture, sniff the content, and select at least four olfactory notes from the list of 22 olfactory notes generated based on the Sigma Aldrich database, which was presented to the participants on a computer screen. Before the series of the odor evaluations, participants were allowed to check the list of olfactory notes and be well aware of the listed olfactory notes. According to the Sigma Aldrich database, some odor components shared the same olfactory notes. (For example, both propan-1-ol and strawberry aldehyde evoked “sweet” notes, while both 4-ethyl-2-methoxyphenol and caryophyllene evoked “meaty” notes.) Therefore, the total number of listed notes was 22 rather than 24. Participants were allowed to provide their own olfactory notes if the smells they perceived were not on the list. Participants were also asked to rank the notes they had provided based on their relative intensity. They were instructed to enter the rankings using a computer. Participants were allowed 30 s to sniff each mixture, and a total of 1 minute to complete the sniffing/evaluation process (Figure 1). After each 1-min evaluation, participants were asked to rest outside the experimental room for 2–3 min to eliminate the effect of residual odors. In total, the psychophysical experiment, where the participant completed the evaluations of the 18 mixtures, lasted approximately 1.5 h.

A similar design was utilized for the odor component experiment, except that the experiment was not repeated, and participants were asked to select/provide at least two olfactory notes.

The olfactory notes and rankings provided by each participant are listed in Supplementary Table 1.

**Figure 1.**
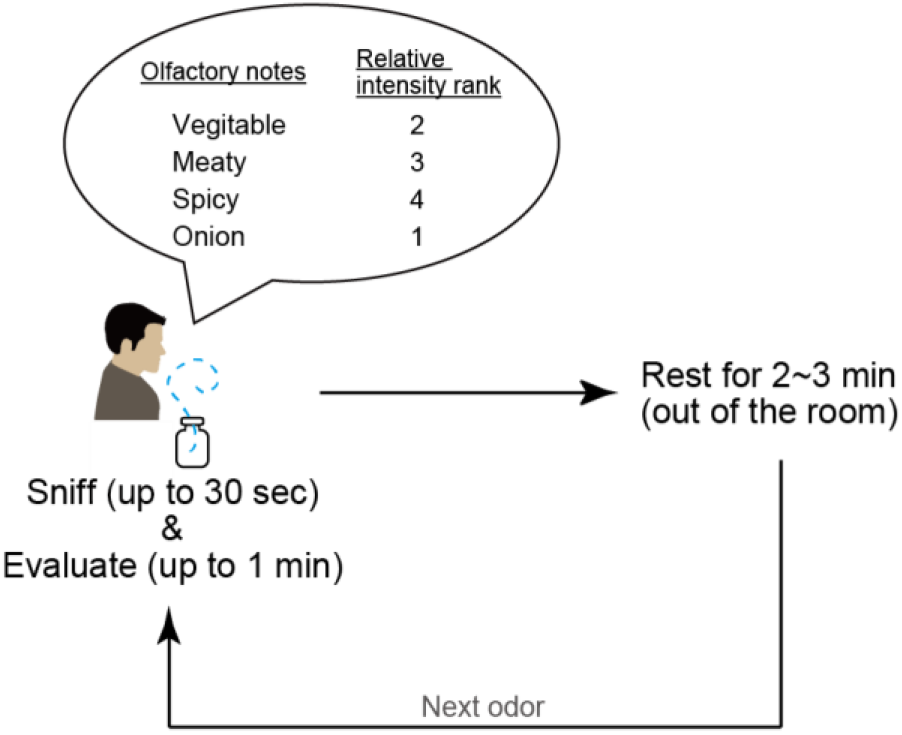
The outline of the psychophysical experiment. Participants sniffed each olfactory stimulus of 18 mixed odors or 12 single odors for up to 30 s, following which they evaluated the olfactory stimulus by selecting olfactory notes and providing a relative intensity ranking for the olfactory stimulus during the remainder of the 1-min period. After the evaluation, participants rested outside of the experimental room, and the process was repeated for a different olfactory stimulus 2-3 min later.

### Data analysis

We defined the relative intensity score (RIS) of an olfactory note as follows:

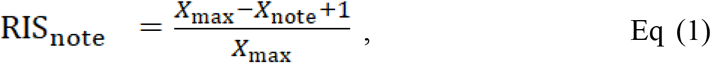

where *note* represents the focal olfactory note, *X*_max_ represents the number of olfactory notes selected by the participant per odor component or mixture, and *X*_note_ represents the rank of the olfactory note provided by the participant based on relative intensity. If the olfactory note was not selected by the participant, the RIS was zero. We then compiled the *m* × *n* RIS data set (*m* = number of the provided notes, *n* = number of participants) for each mixture or component.

Next, we defined the pseudo-mixtures as an idealized mixture in which the olfactory notes of its components had been completely preserved (Figure 2A). The pseudo-mixture was compared to the true mixture evaluated by participants in order to determine whether the olfactory notes of odor components had changed in the mixture. Pseudo-mixture values were calculated as follows: We obtained the matrix 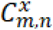 which consisted of the *m* × *n* RIS data set from the result of the odor component experiment, where *x* represents the number of odor samples (1– 12), *m* represents the number of the provided notes by participants (row data), and *n* represents the number of participants (column data). The RIS data sets in 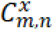 were derived from the results of the component experiment. Using 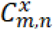, we calculated the RIS-matrix for pseudo-mixture 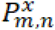 as

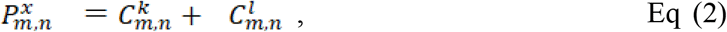

where *k* and *l* represent the number of odor components from 1 to 12 (*k* ≠ *l*). The combination of *x*, *k*, and *l* was based on the data presented in Table 2. For example, pseudo-mixture 1 was regarded as the sum of the RIS-matrices of odor components 1 and 2 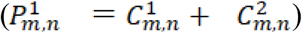, which is illustrated in left of Figure 2A as an example. In contrast to that of the pseudo-mixture, we defined the RIS-matrix of the real mixture based on the actual results of the mixture experiment. Therefore, the RIS data sets for the real and pseudo-mixtures ranged from No. 1 to No. 18, respectively.

**Figure 2.**
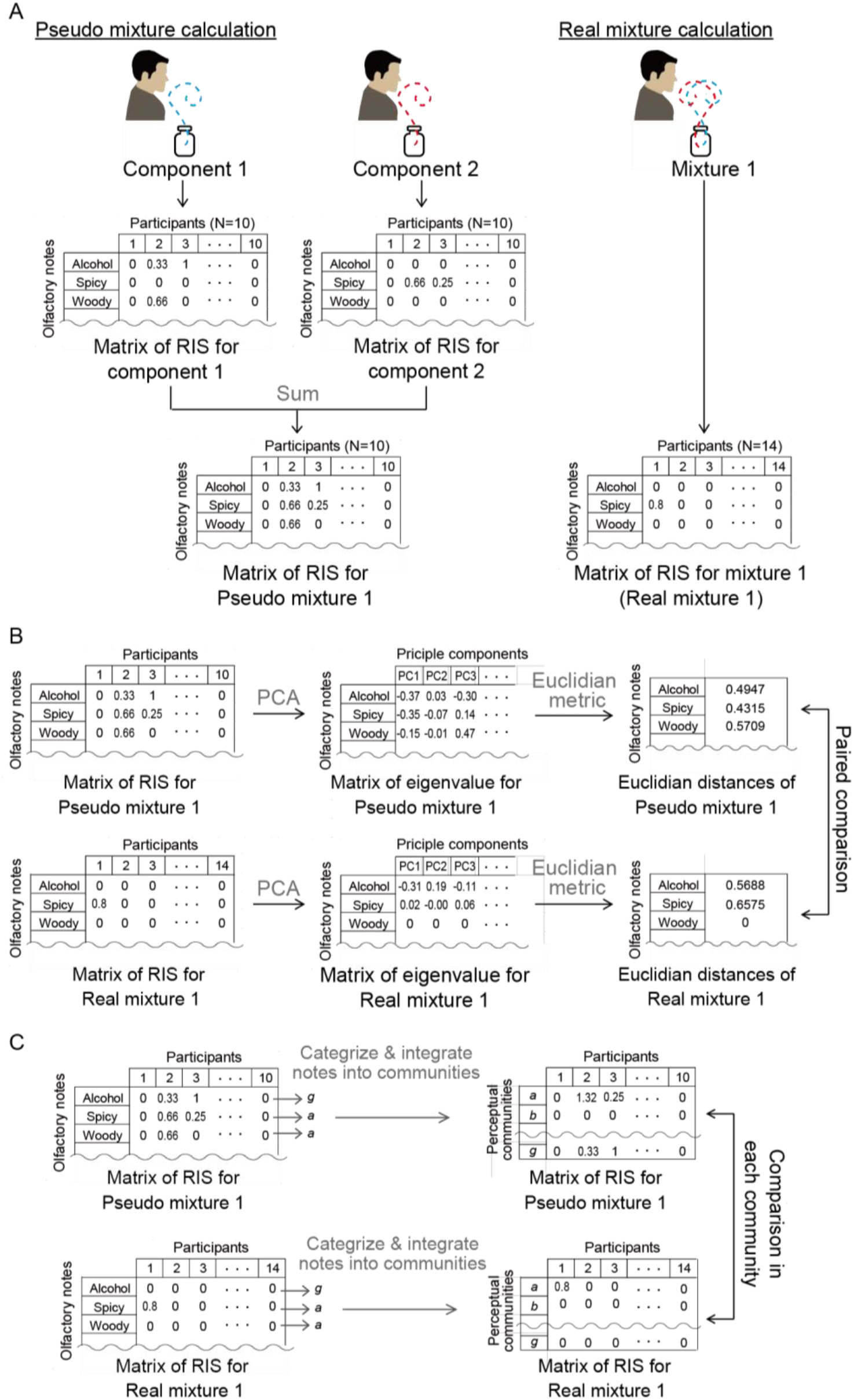
The outline of data analysis. (A) The manner to obtain the data sets of real and pseudo-mixtures is shown. Data sets of the real mixture were derived from the odor mixture experiment and those of pseudo-mixtures were done from the odor component experiment. (B) How to analyze using PCA is shown. Using PCA, we quantified each olfactory note by decreasing the dimensions of participants, and then performed statistical paired comparison between real and pseudo-mixtures using the Euclidian distances. (C) The method of the comparison based on the “perceptual communities” is shown. According to the correspondence described in Table 3, we categorized the olfactory notes into perceptual communities (from *a* to *g*) and performed statistical comparison in each perceptual community.

To evaluate the differences between the real and pseudo-mixtures, we performed comparison based on both individual olfactory notes and major olfactory perceptual groups.

In the comparison of the individual olfactory notes, each olfactory note was quantified using the eigenvalues from principal component analysis (PCA) (Figure 2B). We applied PCA to reduce the dimensions of participants (*n*) and integrated participant responses for each olfactory note. PCA was performed based on the correlation coefficient matrix using JMP 12 (SAS Institute Inc., NC, USA). We then obtained multi-dimensional eigenvalue vectors for each olfactory note. The number of vector dimensions was determined based on the individual cumulative contribution ratio (≥ 80%), respectively. We compared the Euclidean distance of each eigenvalue vector for each pair of real and pseudo-mixtures (e.g., real mixture 1 versus pseudo mixture 1). Notes of “unknown” provided by participants were excluded from statistical analysis.

For comparisons based on major olfactory perceptual groups, we classified the olfactory notes into previously established “perceptual communities”^23^ (Figure 2C). In this previous study, the seven major olfactory perceptual communities (termed communities *a* to *g*) were identified based on odor qualities (*e.g.*, community *a* included plant-related odors such as “herb”, “wood”, etc.). In our study, 90 olfactory notes (22 notes were derived from the Sigma Aldrich database, and 68 notes were provided by participants) were categorized into the seven communities based on the methods of Kumar *et al.* (2015). The degrees of correspondence between the olfactory notes and the communities are listed in Table 3. Non-typeable notes such as “the smell of a hospital” were less than 10% each for real and pseudo-mixture RIS data sets and excluded from the comparison between real and pseudo mixtures. In each pair of real and pseudo-mixtures, the RIS data sets were compared according to perceptual community.

**Table 3.** List of olfactory notes and perceptual communities. The correspondence between olfactory notes used in this study and the “perceptual communities” defined in the previous study of Ritesh Kumar et al. is listed. In the “Community” column, “X” represents the non-typeable notes in the previous study. In the “Derived” column, “S.A.” represents the *Sigma Aldrich Ingredients Catalog: Flavors & Fragrances* and “F.A.” represents the freely answered by participants.

| Olfactory notes | Community | Derived | Olfactory notes | Community | Derived |
| --- | --- | --- | --- | --- | --- |
| Alcohol | g | S. A. | Gelatin | X | F. A. |
| Almond | f | S. A. | Ginger | d | F. A. |
| Apple | c | S. A. | Hay | a | F. A. |
| Cabbage | f | S. A. | Ink | X | F. A. |
| Ethereal | g | S. A. | Insect repellent | e | F. A. |
| Geranium | c | S. A. | Iron | b | F. A. |
| Green | b | S. A. | Energy drink | X | F. A. |
| Hawthorne | c | S. A. | Leaf of marigold | b | F. A. |
| Herbaceous | a | S. A. | Lemon | d | F. A. |
| Honey | c | S. A. | Minty lip balm | X | F. A. |
| Meaty | f | S. A. | Magnolia | c | F. A. |
| Onion | f | S. A. | Manicure | X | F. A. |
| Plum | g | S. A. | Marker pen | X | F. A. |
| Rose | c | S. A. | Felt tip pen | X | F. A. |
| Smoky | a | S. A. | Nail polish remover | X | F. A. |
| Spicy | a | S. A. | Oil | d | F. A. |
| Strawberry | g | S. A. | Ointment | X | F. A. |
| Sulfurous | e | S. A. | Paint | e | F. A. |
| Sweet | g | S. A. | Peach | g | F. A. |
| Vanilla | g | S. A. | Pickle | X | F. A. |
| Vegetable | b | S. A. | Pomegranate | X | F. A. |
| Woody | a | S. A. | Pop | X | F. A. |
| Almond jelly | X | F. A. | Powder medicine with syrup | X | F. A. |
| Ammonia | f | F. A. | Red wine | g | F. A. |
| Antisepsis before injection | f | F. A. | Roast | f | F. A. |
| Baloon | X | F. A. | Rotten | f | F. A. |
| Banana | g | F. A. | Rubber | f | F. A. |
| Bandage | X | F. A. | Seirogan | X | F. A. |
| Burnning garbage | X | F. A. | Similar to Mmixture 17 | X | F. A. |
| Candy | g | F. A. | Smell of grandmather's house | X | F. A. |
| Carrot | b | F. A. | Smell of hospital | X | F. A. |
| Cherry | g | F. A. | Smell of new bag | X | F. A. |
| Cinnamon | a | F. A. | Smell of new shoes | X | F. A. |
| Coarse tea | c | F. A. | Smell of new shop | X | F. A. |
| Coconut | e | F. A. | Smell of roast | f | F. A. |
| Coin | X | F. A. | Smell of rubber boot | f | F. A. |
| Coke | X | F. A. | Sour fruit | X | F. A. |
| Compress | X | F. A. | Sweat | e | F. A. |
| Detergent | X | F. A. | Taping | X | F. A. |
| Earth | f | F. A. | Unknown | X | F. A. |
| Farm | X | F. A. | Varnish | X | F. A. |
| Fish | f | F. A. | Vegetable juice | X | F. A. |
| Food waste | f | F. A. | Vinegar | e | F. A. |
| Gummed tape | X | F. A. | Welsh onion | f | F. A. |
| Garlic | f | F. A. | Yoghurt | e | F. A. |

### Statistics

Because our experimental data did not follow a normal distribution, we used non-parametric tests for statistical analyses.

The Wilcoxon signed-rank test (α = 0.05) was performed for one-by-one comparison of the olfactory notes between the real and pseudo mixtures. We compared the Euclidean distance of each eigenvalue vector of the olfactory note which was obtained by PCA.

The Wilcoxon text were performed to compare data based on perceptual communities (*e.g.*, the RIS score of community *a* was compared between the real and pseudo mixtures). In the comparison of the perceptual communities, Bonferroni α corrections were applied based on the number of the perceptual communities within the pair of real and pseudo mixture. The corrected α levels were as follows: 0.05/7 = 0.0071 in pair of 6 and16 of 1st-day real mixture vs pseudo-mixture; 0.05/6 = 0.0083 in pair of 1, 2, 4, 5, 8, 10, 13, 14, 15, and 18 of 1st-day real mixture vs pseudo-mixture and pair of 1, 2, 4, 5, 6, 12, 14, 17, and 18 of 2nd-day real mixture vs pseudo-mixture; 0.05/5 = 0.01 in pair of 3, 7, 9, 11, 12, and 17 of 1st-day real mixture vs pseudo-mixture and pair of 3, 7, 8, 9, 10, 11, 13, 15, and 16 of 2nd-day real mixture vs pseudo-mixture. Wilcoxon signed-rank tests and Wilcoxon tests were performed using JMP 12 (SAS Institute Inc., NC, USA).

### Gas chromatography/mass spectroscopy analysis

Prior to the psychophysical experiments, we performed gas chromatography and mass spectroscopy (GC/MS) analyses to confirm that the odor components had not undergone any chemical transformations that might affect olfactory perception (Figure 3). GC/MS equipment (7890A GC system, 5975C inert XL MSD with triple-axis detector; Agilent Technologies, Santa Clara, CA, USA) was used to confirm that none of the mixture groups had undergone any chemical reactions. In the sample analyses, each of the four odorant solutions from the same complexity group were spotted onto separate cotton cloths (2 × 2 cm), and the cloths were placed together in a single vial (30 mL). A solid-phase micro extraction fiber (50/30 μm DVB/CAR/PDMS; Sigma Aldrich, St. Louis, MO, USA) was injected into the vial, exposed to the sample headspace for 30 min at 37°C to extract the volatilized and mixed odorants, and the fiber was then immediately transferred to the injection port of the GC/MS machine. A DB-5-ms capillary column (30 mm × 0.25 mm inner diameter with a 0.25-μm film thickness; Agilent Technologies) was used to separate the odorants. The GC oven temperature was increased from 40°C to 130°C at a rate of 3°C/min, and then maintained at 130°C for 30 min to analyze the high-complexity odorants. To analyze the low- and medium-complexity odorants with high volatility, the initial oven temperature was maintained at 40°C for 10 min, increased to 150°C at a rate of 3°C/min, and then to 230°C at 10°C/min, and maintained at a final temperature of 230°C for 20 min. MS was performed in electron ionization mode (70 eV). Chromatogram peaks were identified using the National Institute of Standards and Technology Mass Spectral Library and Aroma Office software (Ver. 5.0; NISHIKAWA KEISUOKU Co., Ltd, Kumamoto, Japan).

**Figure 3.**
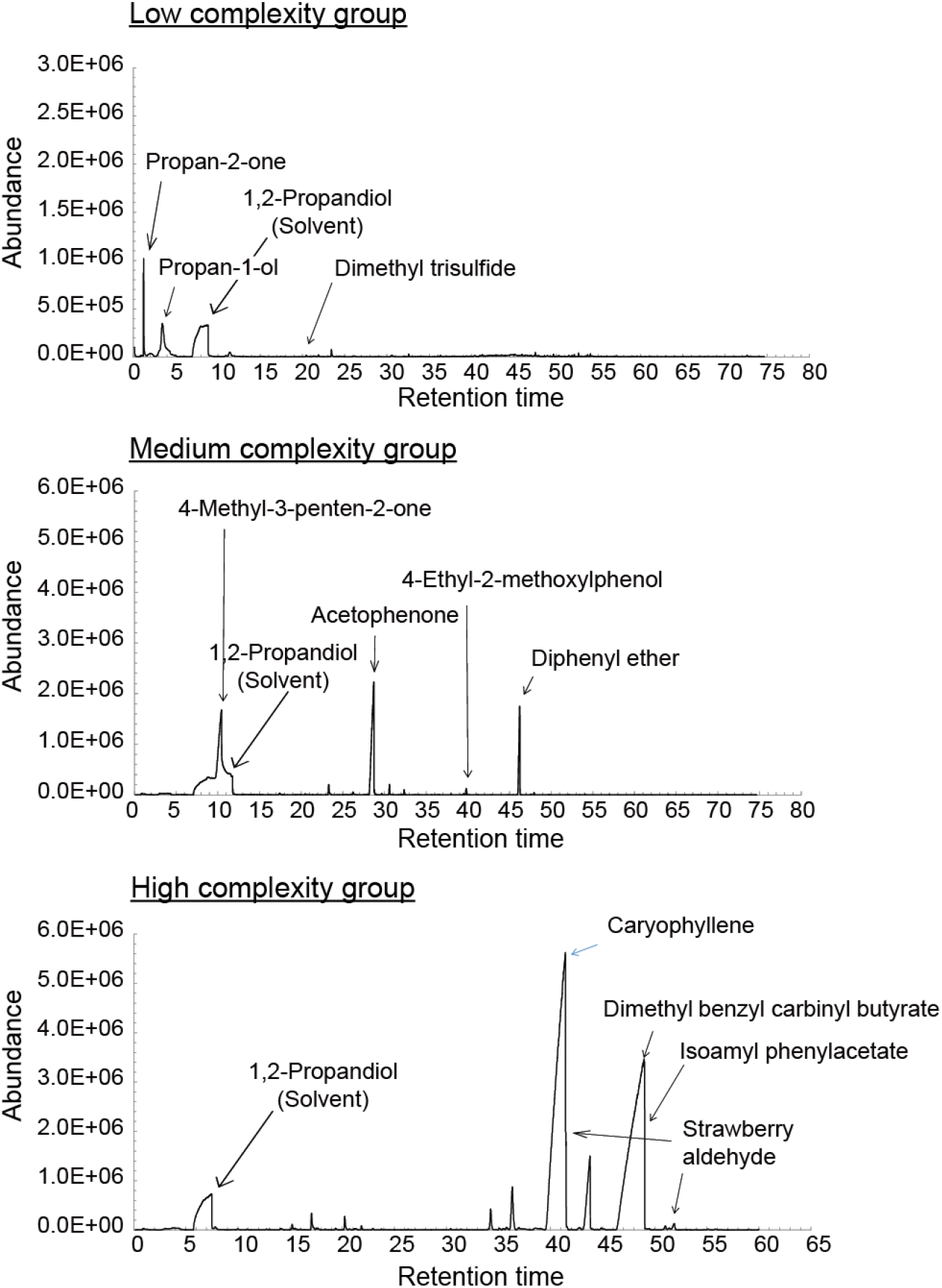
Gas chromatography and mass spectroscopy (GC/MS) analyses for confirmation of no chemical reaction. Chromatographs obtained from the different odorant groups during the GC/MS analyses (top, low-complexity group; middle, medium-complexity group; bottom, high-complexity group). No loss of any of the odor components was noted, and no novel chemical substances were synthesized in the odor mixtures. In the low-complexity group, propane-1-thiol could not be detected due to the extremely low concentration. In the high-complexity group, strawberry aldehyde exhibited dual peaks due to the presence of the optical isomer, and the peak of diastereoisomers was hidden by the peak of dimethyl benzyl carbinyl butyrate.

## Results

### Comparison based on individual olfactory notes

We performed one-by-one comparison of each olfactory note between the real and pseudo-mixtures. The scores of each olfactory note were obtained by PCA (see Experimental and Figure 2A for detail). The analysis revealed that among the 12 mixture pair (1st-day real mixture versus pseudo mixtures and 2nd-day real mixture versus pseudo mixture), low-complexity group showed significance or marginal significance in 8 mixtures, medium-complexity group did in 2 mixtures, and high-complexity group did in 5 mixtures (Wilcoxon signed-rank test, Table 4).

**Table 4.** The result of statistical comparison (Wilcoxon signed-rank test) between real and pseudo mixtures with PCA. The left column shows the number of mixture pair (e.g. real-mixture 1 vs pseudo-mixture 1). The *dfs* were based on the number of quantified olfactory notes. * indicates the significance and † do the marginal significance.

| Complexity group | Mixture pair<br>(real vs pseudo) | Real (1st day) vs<br>Pseudo |  | Real (2ns day) vs<br>Pseudo |  |
| --- | --- | --- | --- | --- | --- |
|  |  | df | p value | df | p value |
| Low complexity | 1 | 23 | 0.0312* | 22 | 0.0885 † |
|  | 2 | 23 | 0.166 | 21 | 0.963 |
|  | 3 | 16 | 0.0092* | 17 | 0.0472* |
|  | 4 | 25 | 0.163 | 24 | 0.0071* |
|  | 5 | 23 | 0.0809 † | 23 | 0.0249* |
|  | 6 | 24 | 0.0864 † | 24 | 0.274 |
| Medium complexity | 7 | 30 | 0.238 | 26 | 0.542 |
|  | 8 | 25 | 0.256 | 24 | 0.556 |
|  | 9 | 24 | 0.179 | 26 | 0.0014* |
|  | 10 | 29 | 0.594 | 28 | 0.452 |
|  | 11 | 25 | 0.912 | 25 | 0.630 |
|  | 12 | 23 | 0.300 | 26 | 0.0338* |
| High complexity | 13 | 24 | 0.0002* | 22 | 0.077 † |
|  | 14 | 19 | 0.151 | 23 | 0.307 |
|  | 15 | 19 | 0.748 | 27 | 0.517 |
|  | 16 | 22 | 0.649 | 22 | 0.626 |
|  | 17 | 23 | 0.0497* | 28 | 0.353 |
|  | 18 | 20 | 0.0703 † | 21 | 0.095 † |

We found that more number of the low-complexity mixtures showed the significance and marginal significance than medium- and high-complexity group in the comparison of real and pseudo mixtures.

### Comparison based on major perceptual group

We compared the real and pseudo-mixtures by major olfactory perceptual groups, “perceptual community”^23^. We classified the olfactory notes into the perceptual communities according to the previous study, then obtained the RIS for each community in each mixture. The comparison by the communities showed that among the 12 mixture pair (1st-day real mixture versus pseudo mixtures and 2nd-day real mixture versus pseudo mixture), low-complexity group showed significance or marginal significance in 7 mixture pairs, medium-complexity group did in 2 mixture pairs, and high-complexity group did in 2 mixture pairs (Wilcoxon rank-sum test with Bonferroni α correction, Table 5 and 6).

**Table 5.**
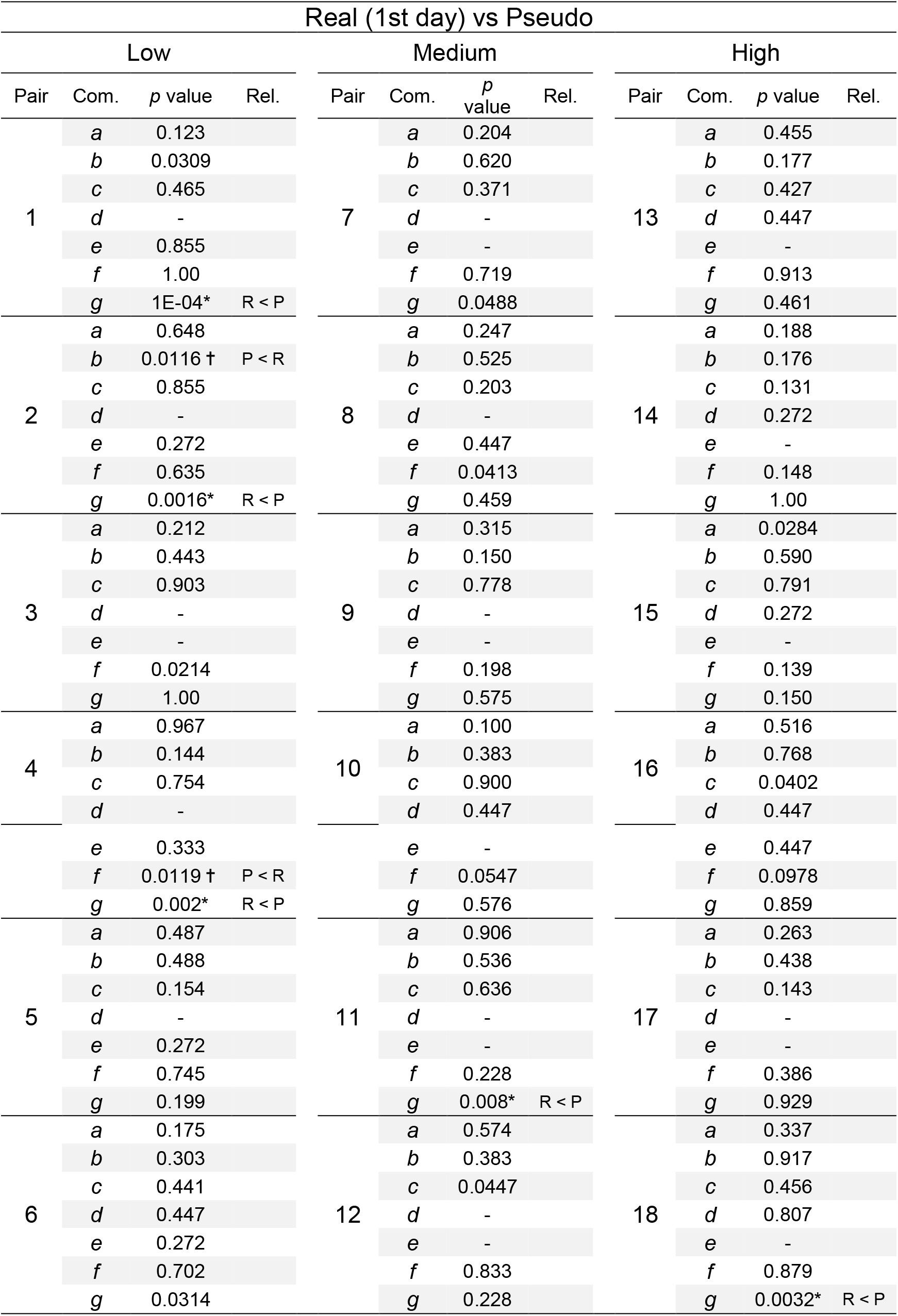
The result of statistical comparison by the perceptual community between real (1st day) and pseudo mixtures. The column of “Pair” indicates the number of mixture pair (*e.g.*, real-mixture 1 vs pseudo-mixture 1). The column of “Com.” indicates the perceptual community. Wilcoxon rank-sum tests were performed to compare relative intensity scores between real mixtures (obtained from 14 participants of the mixture experiment and pseudo-mixtures (obtained from 10 participants of the component experiment). The α level was Bonferroni corrected. Dashes indicate “0” responses, which were excluded when Bonferroni correction was performed. In the “Rel.” column, “Rel.” represents the relation of the relative intensity scores between each pair of real and pseudo mixture, “R” refers to the real mixture, while “P” refers to the pseudo-mixture. The relation was shown only in the perceptual community exhibiting significant difference. * indicates the significance and † do the marginal significance.

**Table 6.**
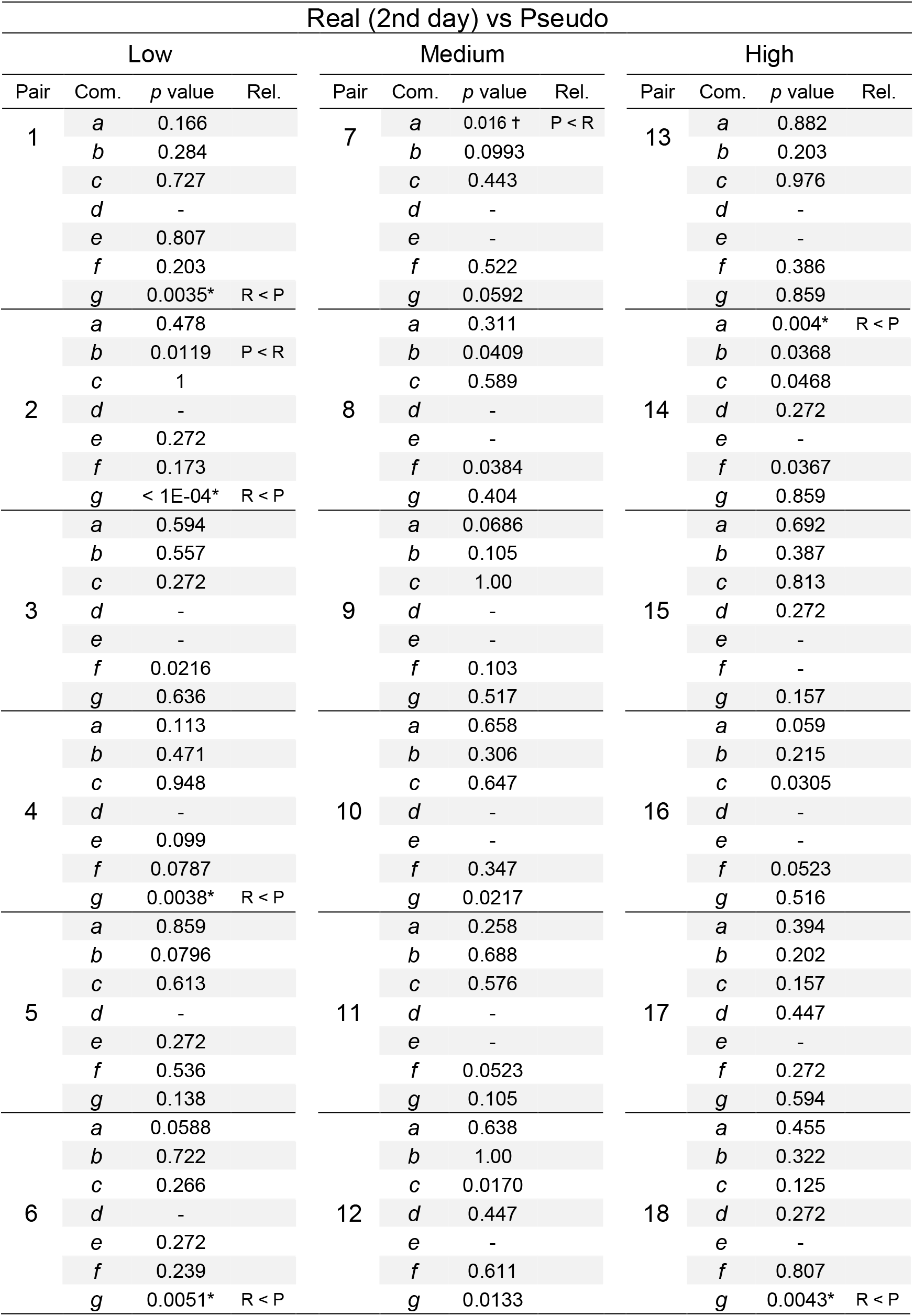
The result of statistical comparison by the perceptual community between real (2nd day) and pseudo mixtures. The column of “Pair” indicates the number of mixture pair (*e.g.*, real-mixture 1 vs pseudo-mixture 1). The column of “Com.” indicates the perceptual community. Wilcoxon rank-sum tests were performed to compare relative intensity scores between real mixtures (obtained from 14 participants of the mixture experiment) and pseudo-mixtures (obtained from 10 participants of the component experiment). The α level was Bonferroni corrected. Dashes indicate “0” responses, which were excluded when Bonferroni correction was performed. In the “Rel.” column, “Rel.” represents the relation of the relative intensity scores between each pair of real and pseudo mixture, “R” refers to the real mixture, while “P” refers to the pseudo-mixture. The relation was shown only in the perceptual community exhibiting significant difference. * indicates the significance and † do the marginal significance.

The results exhibited that more number of the low-complexity mixtures showed the significance and marginal significance than medium- and high-complexity group in the comparison of real and pseudo mixtures.

## Discussion

In the present study, we demonstrated that odor mixtures composed of low-complexity odorants were perceived as relatively different smells from those of the original components. To evaluate whether the smells of odor components had changed when mixed with other components, we compared participant responses to a series of pseudo-mixtures, which were presumed to retain the olfactory notes of the individual components. PCA of individual olfactory notes revealed that real low-complexity mixtures induced different olfactory notes than those indicated for pseudo-mixtures. Analysis of olfactory notes based on seven major perceptual communities^23^ revealed that real and pseudo-mixtures of the low-complexity group differed significantly with regard to perceptual community. Thus, our findings can suggest that humans have the olfactory capacity to detect a specific smell from among a mixture, depending on the complexity of its odor components.

In the psychological experiments, the two analyses conducted in the present study exhibited consistent results, in which low-complexity odorants were relatively perceived as novel smells when individual components were mixed with one another. We examined differences between real and pseudo-mixtures by comparing data among individual olfactory notes or perceptual communities. Low-complexity mixtures 1, 4, and 6 exhibited significant or marginally significant differences in both comparisons of individual olfactory notes and those based on perceptual community (Table 4, 5, and 6). Our results suggest that participants perceived the smells of these mixtures as distinct from their components with regard to both specific and general odor qualities. For other low-complexity mixtures, significant differences in either olfactory notes or perceptual community were observed. In mixtures 3 and 5, we observed significant differences only in the comparison of olfactory notes. Such results indicate that the real and pseudo-mixtures were perceived as similar in quality, although participants were capable of differentiating the smells verbally. In the present study, participants carefully evaluated each smell by referring to a pre-determined list of olfactory notes. Furthermore, the number of olfactory notes selected from the list was much higher than the number of terms provided by participants (ratio of listed to provided notes = 13:3 in 1st-day real mixture 3; 12:5 in 2nd-day real mixture 3; 10:0 in pseudo mixture 3; 18:2 in 1st-day mixture 5; 18:2 in 2nd-day mixture 5; 14:3 in pseudo mixture 5). These results indicate that minor alterations in olfactory notes may have occurred within the perceptual community of mixtures 3 and 5. For mixture 2, significant differences were observed only in the analysis of perceptual communities. This result may be attributed to the property of the olfactory verbalization. Previous studies have demonstrated that the verbalization of aspects related to olfactory stimuli is more difficult than that for other senses such as vision^25^. Thus, limitations in olfactory verbalization may result in the perception of real and pseudo-mixtures as qualitatively different, even if the difference cannot be verbalized when presented with a list of options. In contrast to findings observed for low-complexity mixtures, few medium- or high-complexity mixtures exhibited significant differences with regard to olfactory notes or perceptual community. Thus, these findings indicate that the smells of low-complexity mixtures are perceived as different from those of their components with regard to specific (olfactory verbal expression) and/or general qualities (perceptual community).

We next discuss the validity of the methods used to analyze differences among individual olfactory notes and perceptual communities. PCA was applied to quantitatively evaluate data for each olfactory note. In previous studies, PCA was utilized to identify the major olfactory perceptual groups or characterize odor profiles by reducing the dimensions of olfactory perceptual descriptors^19,30,31^. In the present study, we utilized PCA to quantify each olfactory note by reducing the dimensions of participant responses, which enabled us to perform the statistical comparison between the real and pseudo-mixtures. By comparing individual olfactory notes, we aimed to increase the sensitivity of detecting alterations in olfactory perception, even if the alteration was minor (*e.g.*, transformation from “Rose” to “Geranium”). In our subsequent analyses, we compared differences in perceptual communities between the real and pseudo-mixtures. The seven perceptual communities were established in a previous study by performing a network analysis of numerus olfactory notes obtained from several databases, including *Sigma Aldrich Ingredients Catalog: Flavors & Fragrances*^23^. By comparing the communities, we intended to examine whether general odor quality differed substantially between the real and pseudo-mixtures. Thus, the use of both analyses enabled us to detect both minor/specific and substantial/general alterations in olfactory perception between the real and pseudo-mixtures.

The selection of olfactory notes can be affected by a participant’s lexical knowledge^32^. In the present study, we limited the number of olfactory notes and instructed participants to select notes from among those on an existing list, enabling us to control for differences in the lexical background of participants. A previous study reported that single, high-complexity odorants induced more notes^21^. This previous study focused on the number of olfactory notes, and participants were instructed to freely provide their own olfactory notes. In contrast, our study focused on the olfactory notes themselves rather than on the number of notes, and participants were instructed to select at least four (odor mixture experiment) or two (odor component experiment) olfactory notes from among those on the given list in the component or mixture experiment, respectively. The selection of olfactory notes from the list would drive the participants towards elemental perception at the expense of configural perception^33^. However, we observed that low-complexity odor mixtures exhibited differences in both olfactory notes and perceptual quality between real and pseudo-mixtures. Furthermore, our analyses confirmed that the total number of olfactory notes provided by the participants did not significantly differ among the odor component and mixtures used in the present study (Supplementary Figure S1). Thus, our findings indicate that molecular complexity plays a role in determining the perceptual mode, and that the findings were not affected differences in the number of olfactory notes provided.

The present study has several limitations. First, the number of odorants and mixtures investigated in our experiments may have been insufficient for deriving a definitive conclusion. To examine the association between molecular complexity and perceptual mode, we utilized odor components and mixtures with a discrete rather than continuous range of complexity scores. Although differences between real and pseudo-mixtures were observed in the low-complexity group, we cannot exclude the possibility that other ranges of complexity scores (*e.g.*, 50–100) would yield different results. Furthermore, in our analysis of perceptual communities, the insufficient variety of odorants may explain why significant differences were observed only for community *g* in the low-complexity group. Comprehensive investigations using a greater number of odorants and mixtures may enable researchers to analyze the perceptual profiles of individual mixtures in detail. Second, we simplified the design of our study by including only binary odor mixtures, although odor mixtures in the real world are often composed of numerous odorants. Therefore, our finding that low-complexity mixtures induce configural perception may be restricted to binary odor mixtures. Although our findings partially elucidate the association between molecular features of odorant molecules and olfactory perception, future studies should examine this association for mixtures of three or more odorants to improve the generalizability of our findings. Third, the neural mechanism how the molecular complexity influenced the odor mixture perception remains to be concluded from our results. A previous study suggested that low-complexity odorants activated the less number of olfactory receptors to elicit the less variety olfactory notes^21^, whereas nonlinear olfactory processing can be performed for mixture perception^28,29^. To elucidate the relationship between the complexity and mixture perception, neural live-imaging of human olfactory receptor and bulb needs to be established.

In conclusion, the findings of the present study can suggest that molecular complexity influences the olfactory perceptual mode of odor mixtures. We observed that odor mixtures composed of low-complexity odorants were perceived as relatively novel odors, indicating that molecular complexity may influence how the odorant and receptor interact to produce the associated neural representation in the central olfactory system. Such information may further our understanding of the olfactory perceptual modes of odor mixtures.

## Acknowledgements

We would like to thank members of our laboratory for the discussions.

Funding: This work was supported by Kobayashi International Scholarship Foundation and Qdai-jump Research Program [grant number 27818, 2015-2017].

## Authors and Contributors

MH and TO designed the study. MH performed the psychological experiments. HI and YK performed the GC/MS analyses. MH, KT, and TO analyzed the data. MH wrote the manuscript. TO, KT, YK, and HI reviewed the manuscript. TO and HI supervised the study.

## Data availability

The data that support the findings of this study are available from the corresponding author upon reasonable request.

## Supplementary materials

**Supplementary Figure S1.**
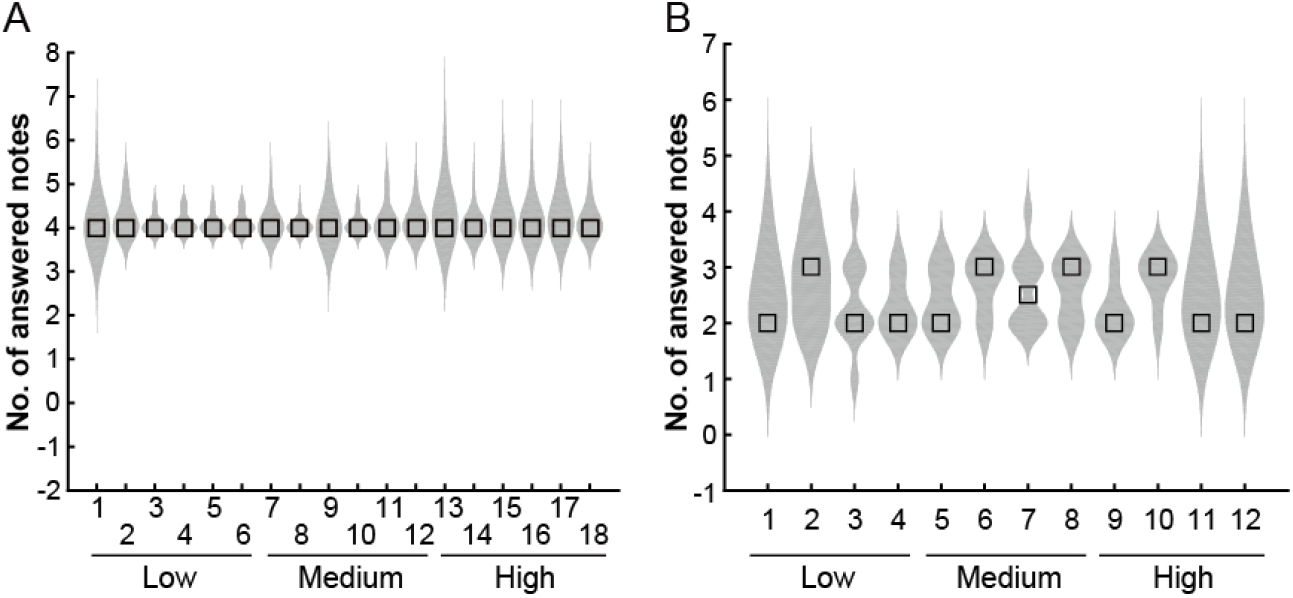
The number of provided olfactory notes. The y-axis shows the total number of olfactory notes provided per mixture (A) or component (B). Wilcoxon tests revealed no significant differences among the mixtures or components (A: *p* = 0.71, B: *p* = 0.58).

**Supplementary Table 1 Law data of participant responses** (see other file)

Blank spaces indicate that a participant did not record a response for relative intensity, and the blanks were excluded from the analyses. RIR means the relative intensity rankings.

## References

[1] S. Barkat, E. Le Berre, G. Coureaud, G. Sicard, T. Thomas-Danguin, Chem. Senses 2012, 37, 159–166.

[2] J. D. Howard, J. A. Gottfried, Neuron 2014, 84, 857–869.

[3] D. G. Laing, G. W. Francis, Physiol. Behav. 1989, 46, 809–814.

[4] N. Y. Schneider, F. Datiche, D. A. Wilson, V. Gigot, T. Thomas-Danguin, G. Ferreira, G. Coureaud, Brain Struct. Funct. 2015, DOI 10.1007/s00429-015-1055-2.

[5] T. Thomas-Danguin, C. Sinding, S. Romagny, F. El Mountassir, B. Atanasova, E. Le Berre, A.-M. Le Bon, G. Coureaud, Front. Psychol. 2014, 5, 504.

[6] A. Jinks, D. G. Laing, Physiol. Behav. 2001, 72, 51–63.

[7] K. Marshall, D. G. Laing, A. L. Jinks, I. Hutchinson, Chem. Senses 2006, 31, 539–545.

[8] A. Livermore, D. G. Laing, Percept. Psychophys. 1998, 60, 650–61.

[9] A. Livermore, D. G. Laing, J. Exp. Psychol. Hum. Percept. Perform. 1996, 22, 267–77.

[10] S. Romagny, G. Coureaud, T. Thomas-Danguin, Flavour Fragr. J. 2017, 97–105.

[11] L. Buck, R. A. Here, S. Firestein, C. Greer, P. Mombaerts, R. Axel, Cell 1991, 65, 175–187.

[12] Y. Oka, S. Katada, M. Omura, M. Suwa, Y. Yoshihara, K. Touhara, Neuron 2006, 52, 857–869.

[13] K. M. Igarashi, K. Mori, J. Neurophysiol. 2004, 1007–1019.

[14] J. D. Howard, J. Plailly, M. Grueschow, J.-D. Haynes, J. A. Gottfried, Nat. Neurosci. 2009, 12, 932–8.

[15] K. Mori, Y. K. Takahashi, K. M. E. I. M. Igarashi, M. Yamaguchi, Physiol. Rev. 2006, 86, 409–33.

[16] Y. K. Takahashi, M. Kurosaki, S. Hirono, K. Mori, J. Neurophysiol. 2004, 92, 2413–2427.

[17] N. Uchida, Y. K. Takahashi, M. Tanifuji, K. Mori, Nat. Neurosci. 2000, 3, 1035–1043.

[18] B. A. Johnson, C. C. Woo, M. Leon, J. Comp. Neurol. 1998, 393, 457–471.

[19] R. M. Khan, C. Luk, A. Flinker, A. Aggarwal, H. Lapid, R. Haddad, N. Sobel, J. Neurosci. 2007, 27, 10015–10023.

[20] J. Hendrickson, P. Huang, A. G. Tozcko, J. Chem. Inf. Comput. Sci. 1987, 27, 63–67.

[21] F. Kermen, A. Chakirian, C. Sezille, P. Joussain, G. Le Goff, A. Ziessel, M. Chastrette, N. Mandairon, A. Didier, C. Rouby, M. Bensafi, Sci. Rep. 2011, 1, 1–6.

[22] A. Keller, L. B. Vosshall, bioRxiv 2016, 49999.

[23] R. Kumar, R. Kaur, B. Auffarth, A. P. Bhondekar, PLoS One 2015, 10, 1–19.

[24] C. Bushdid, M. O. Magnasco, L. B. Vosshall, A. Keller, Science (80-.). 2014, 343, 1370–1372.

[25] J. K. Olofsson, J. A. Gottfried, Trends Cogn. Sci. 2015, 19, 314–321.

[26] Y. Furudono, Y. Sone, K. Takizawa, J. Hirono, T. Sato, Chem. Senses 2009, 34, 151–158.

[27] L. M. Kay, C. A. Lowry, H. A. Jacobs, Behav. Neurosci. 2003, 117, 1108–14.

[28] D. E. Frederick, L. Barlas, A. Ievins, L. M. Kay, Behav. Neurosci. 2009, 123, 430–437.

[29] K. J. Grossman, A. K. Mallik, J. Ross, L. M. Kay, N. P. Issa, Eur. J. Neurosci. 2008, 27, 2676–2685.

[30] A. A. Koulakov, Front. Syst. Neurosci. 2011, 5, 1–8.

[31] M. Zarzo, D. T. Stanton, Chem. Senses 2006, 31, 713–724.

[32] J. Poncelet, F. Rinck, F. Bourgeat, B. Schaal, C. Rouby, M. Bensafi, T. Hummel, Behav. Brain Res. 2010, 208, 458–465.

[33] E. Le Berre, T. Thomas-Danguin, N. Béno, G. Coureaud, P. Etiévant, J. Prescott, Chem. Senses 2008, 33, 193–199.

